# spatiAlign: An Unsupervised Contrastive Learning Model for Data Integration of Spatially Resolved Transcriptomics

**DOI:** 10.1101/2023.08.08.552402

**Authors:** Chao Zhang, Lin Liu, Ying Zhang, Mei Li, Shuangsang Fang, Qiang Kang, Ao Chen, Xun Xu, Yong Zhang, Yuxiang Li

**Author notes:** These authors contributed equally to this work (Chao Zhang, Lin Liu, Ying Zhang).

## Abstract

Integrative analysis of spatially resolved transcriptomics datasets empowers a deeper understanding of complex biological systems. However, integrating multiple tissue sections presents challenges for batch effect removal, particularly when the sections are measured by various technologies or collected at different times. Here, we propose spatiAlign, an unsupervised contrastive learning model that employs the expression of all measured genes and the spatial location of cells, to integrate multiple tissue sections. It enables the joint downstream analysis of multiple datasets not only in low-dimensional embeddings but also in the reconstructed full expression space. In benchmarking analysis, spatiAlign outperforms state-of-the-art methods in learning joint and discriminative representations for tissue sections, each potentially characterized by complex batch effects or distinct biological characteristics. Furthermore, we demonstrate the benefits of spatiAlign for the integrative analysis of time-series brain sections, including spatial clustering, differential expression analysis, and particularly trajectory inference that requires a corrected gene expression matrix.

## Introduction

The rapid advancements of spatially resolved transcriptomics (SRT) have revolutionized our understanding of the spatial organization and heterogeneity of cells within complex tissues and developmental processes^1^. Cutting-edge in situ capturing technologies (e.g., 10x Genomics Visium^2^, Slide-seq^3^, Stereo-seq^4^, and Seq-scope^5^) have facilitated the simultaneous measurement of tens of thousands of genes in their spatial context, achieving unprecedented cellular or even subcellular resolution. The SRT datasets are typically acquired from different tissue sections, each potentially representing a fragmented profiling of the targeted biological system. Hence, integrating multiple datasets for joint analysis is imperative to decipher the whole biological system. However, integrative analysis presents significant challenges due to the inherent biological variability and batch effects caused by nonbiological factors such as technology differences and different experimental batches.

Prior efforts to tackle this task have conventionally focused on single-cell RNA sequencing technologies (scRNA-seq)^6, 7^, which can be roughly classified into two main categories: methods that (1) generate a joint embedding space^8-13^ and (2) calculate a corrected feature matrix^14-17^. For example, Harmony^8^ projects cells into a shared embedding by maximum diversity clustering and iteratively learning a cell-specific linear correction function that regresses out biological effects within clusters. SCALEX^13^, a deep learning method, provides a truly online tool to project cells into a batch-invariant, common cell-embedding space. Although these methods prove valuable for capturing the overall characteristics of cells, such as combined clustering, they are not applicable to downstream gene-level analysis tasks, such as differentially expressed gene (DEG) analysis. In contrast, popular MNN-based methods such as Seurat v3^16^ efficiently address batch effects in gene expression, but their limitation lies in the ability to align only two batches at a time, and they become impractical when dealing with many batches. However, it is worth noting that these scRNA-seq data integration tools have focused on harmonizing gene expression profiles across different experimental batches and do not consider the spatial context of spots/cells.

In the field of SRT studies, embedding spatial information has proven beneficial for downstream analysis, such as spatial domain identification^18, 19^, imputation^20, 21^, clustering^22^, and cell-type annotation^23^. More recently, works have been published to improve the integration of SRT datasets by exploiting spatial information. PRECAST leverages spatial smoothness in both the cluster label and lower-dimensional representations to estimate aligned embeddings for multiple tissue sections, effectively capturing the spatial relationship between cells/spots^24^. GraphST introduces a graph self-supervised contrastive learning model to reconstruct gene expression by minimizing the embedding distance between spatially adjacent spots^25^. However, PRECAST only returns the corrected embedding space, and GraphST requires registering the spatial coordinates of samples first to ensure its integration performance; thus, their applications are limited in certain scenarios.

To address these challenges, we propose spatiAlign, an unsupervised method that leverages spatial embedding and across-domain adaptation strategies for aligning SRT datasets. spatiAlign offers three key advantages as follows. First, it effectively captures the underlying relationships between spots/cells in both the spatial neighbourhoods and gene expression to learn latent representations with a deep graph infomax (DGI)^26^ framework. Second, spatiAlign aligns biological effects by adapting the semantic similarities between spots/cells and/or pseudoclusters from one section to another without relying on external labelled data, resulting in a joint batch-corrected embedding. Third, benefiting from a symmetric decoder in DGI, spatiAlign outputs the reconstructed spatial gene expression matrices, in which gene expression is enhanced and batch effects are corrected. We validate the three advantages of spatiAlign with four applications on publicly available 10x Genomics Visium, Slide-seq, and Stereo-seq datasets of human and mouse tissues. The benchmarking analysis demonstrates spatiAlign’s superiority in learning low-dimensional representations compared with eight established methods, including GraphST and PRECAST, which were recently developed for SRT datasets. Compared with the original spatial expression of brain region-specific markers, the reconstructed counts from spatiAlign better reflect their laminar organization with denoised, enhanced expressions and clear boundaries between regions. We also validate the capability of spatiAlign to capture the unique characteristics of three Slide-seq mouse hippocampus slices, which contain regions with different structures. The comprehensive integrated analysis of developing moues brain slices indicates that the aligned joint representations, which embed cellular neighbourhoods, improve the identification of cell clusters. In addition, the reconstructed features from our proposed spatiAlign method facilitate the identification of DEGs under different developmental stages and the recovery of cellular trajectories.

## Results

### Overview of spatiAlign

spatiAlign takes as inputs multiple SRT datasets, comprising the expression of all measured genes and spatial locations of spots/cells, to achieve two objectives: low-dimensional semantic alignment and high-dimensional gene expression reconstruction (Fig. 1a). In low-dimensional alignment, the primary strategy underlying spatiAlign is to implement a self-supervised contrastive learning architecture (DGI-based framework) for dimensional reduction while simultaneously propagating neighbouring spatial context between spots/cells (Fig. 1c). Furthermore, it employs an across-domain adaptation technique to align joint embeddings, effectively accounting for batch effects across multiple tissue sections (Fig. 1b). In high-dimensional gene expression reconstruction, we utilize a decoder included in the DGI to reverse aligned representations back into the raw gene expression space, thereby enhancing the gene expression counts.

**Fig. 1.**
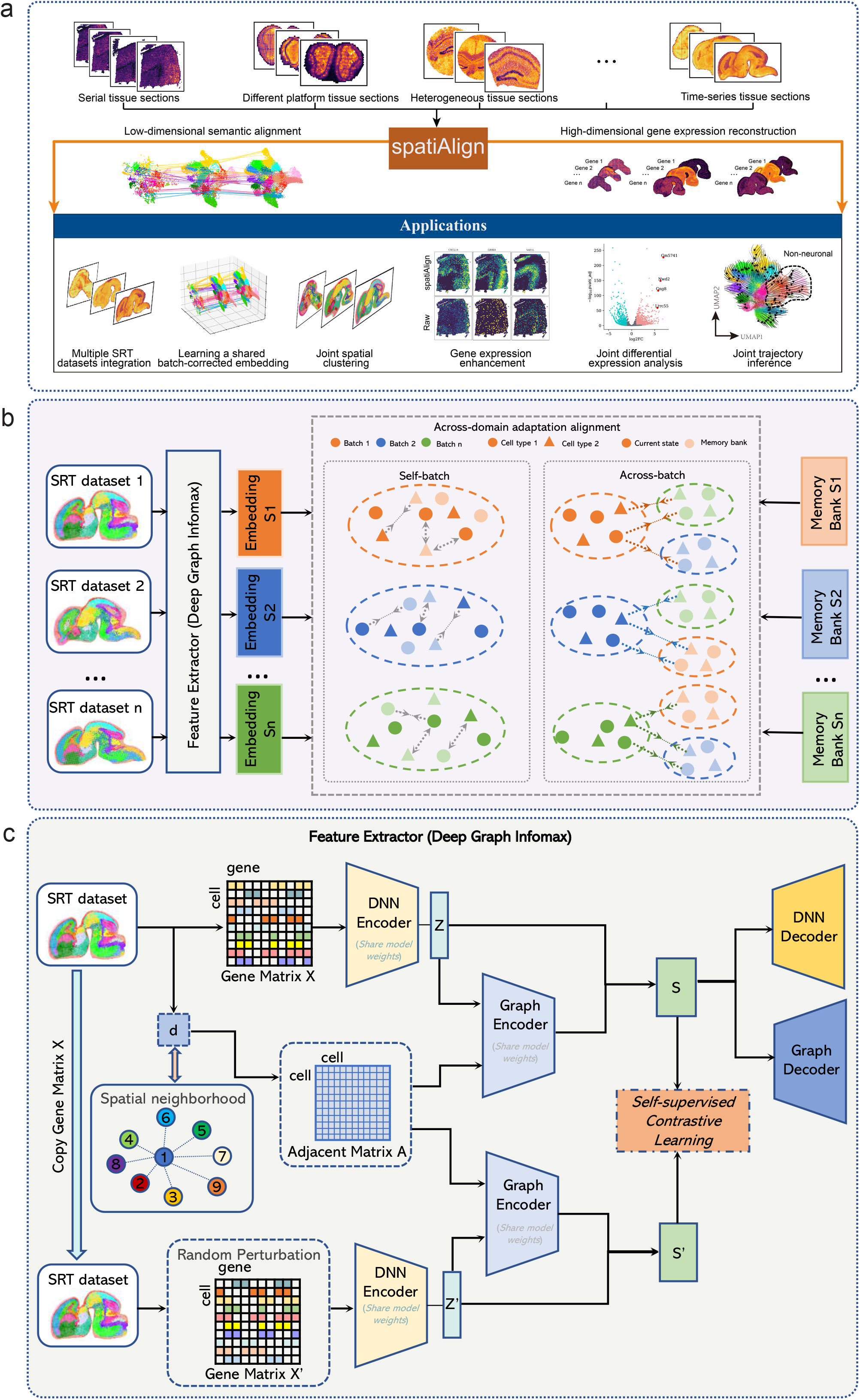
Overview of spatiAlign. **a)**. spatiAlign takes as inputs multiple spatially resolved transcriptomics (SRT) datasets that consist of gene expression profiles for all measured genes and spatial locations of spots/cells. Using semantic alignment, spatiAlign generates a shared batch-corrected embedding, where biological effects are aligned. Moreover, spatiAlign reconstructs the full high-dimensional expression space, enhancing and correcting gene expression counts. In addition to SRT dataset integration and gene feature correction, spatiAlign returns a final joint embedding and enhanced gene expression matrices to facilitate downstream analysis, such as joint spatial clustering, joint differential expression analysis, and joint trajectory inference. **b)**. spatiAlign takes multiple SRT datasets as inputs. Latent embeddings are first generated using Deep Graph Infomax (DGI) as feature extractors. Then, with the utilization of across-domain adaptation and memory bank strategies, sptiAlign brings similar semantic spots/cells closer together and pushes dissimilar spots/cells farther apart, irrespective of their original datasets. These self-batch and across-batch contrastive learning processes align biological effects while correcting batch effects. **c)**. A DGI framework takes as inputs the normalized gene expression matrix and corresponding spatial coordinates from an SRT dataset. A spatial neighbouring graph (i.e., adjacent matrix *A*) is built to represent the spatial relationships between adjacent spots/cells. To create an augmented gene expression matrix *X*′, a random perturbation is applied to shuffle the original gene expression *X* while maintaining the spatial neighbouring graph unchanged. Deep neural network (DNN)-based autoencoders are used to learn gene representations *Z* and *Z*′ by reducing the dimension of gene expression matrix *X* and the augmented expression matrix *X*′. These representations are individually fed into a variational graph autoencoder (VGAE), along with the spatial neighbouring graph, which performs spatial embedding for the gene representations and outputs the final latent representations *S* and *S*′ that capture the rich information both in original/augmented gene expression profiles and spatial information. Afterwards, embeddings *S* are optimized using our self-supervised contrastive learning strategy, which ensures that spatially adjacent cells have similar embeddings while nonadjacent cells have dissimilar embeddings. Finally, the final embeddings *S* can be reversed back to the original feature space, resulting in a reconstructed gene expression matrix.

Formally, given a series of SRT datasets, gene expression profiles are transformed into cell/spot-gene matrices (e.g., gene expression matrix *X*) and spatial neighbouring graphs between cells/spots (e.g., cell‒cell adjacent matrix *A*), where the connective relationships of cells/spots are negatively associated with Euclidean distance. We design a deep neural network (DNN)-based autoencoder to learn the low-dimensional gene representations *Z* from the original gene expression matrix. The adjacency matrix *A* and the reduced gene representations *Z* are fed into a variational graph autoencoder (VGAE)^27^ that propagates spatial neighbouring context for the gene representations, resulting in a final joint representation *S* (positive samples) that captures comprehensive characteristics of the gene expression profile and cellular neighbourhoods. Thereafter, the enhanced gene expression matrices can be reconstructed using a symmetric decoder architecture, which reverses the joint representations *S* back to the original space.

To improve spatiAlign’s ability to exploit potential information in SRT datasets, augmentation-based contrastive learning is adopted^25, 28, 29^. Technically, a gene expression matrix *X* is augmented by randomly shuffling the gene expression vector of spots/cells to create a corrupted gene expression matrix *X* while keeping the spatial neighbouring graph unchanged. Then, the corrupted gene expression matrix *X* and adjacency matrix *A* are fed into the aforementioned model, which utilizes the shared model weights to generate corrupted joint representations *S* (negative samples). We then use self-supervised contrastive learning to bring the positive samples closer within the spatial neighbouring context while pushing the negative samples far apart within the same neighbouring context (Fig. 1c).

Using an across-domain adaptation^28, 30, 31^ and deep clustering^32^ strategy, spatiAlign aims to align biological effects while maximizing the preservation of biological variances in the latent embedding of spots/cells. Specifically, we use a memory bank to store the final latent representations for each dataset that will be used to measure the similarity between spots/cells or pseudoclusters for self-batch/across-batch contrastive learning. For each tissue section, spatiAlign minimizes the similarity distance between the current latent representations and the corresponding memory bank entries to bring similar semantic spots/cells closer together and push dissimilar semantic spots/cells far apart. In parallel, inspired by the idea of “label as representation”, we assume that the dimension of the final latent embedding is equal to the number of pseudoprototypical clusters, and the spots/cells vector denotes its soft label accordingly. Thus, each spot/cell is assigned to a different pseudo cluster, and all pseudo clusters should differ from each other. Identically, spatiAlign employs “current pseudocluster representation” (transposition latent representation) and “cached pseudocluster representation” (transposition corresponding memory bank) to bring the same pseudocluster spots/cells closer together and push dissimilar pseudo cluster spots/cells far apart, avoiding pseudocluster dropout intrinsic biological variances. In across-batch contrastive learning, cross-similarity between spots/cells, measured by the current latent representation and memory bank of other sections, is minimized to align biological effects across sections, ensuring similar semantic spots/cells closer together, regardless of which sections they are from.

### spatiAlign outperforms the control methods in integrating DLPFC datasets

We evaluated the effectiveness of spatiAlign in analysing a series of 10x Genomics Visium datasets from the human dorsolateral prefrontal cortex (DLPFC). The selected dataset comprised four sections that were manually annotated into six tissue layers (Layer_1 to Layer_6) and white matter (WM) in the original study (Fig. 2a, Supplementary Fig. S1a)^33^. We first performed graph-based clustering (Leiden) on the latent representations of spatiAlign and the other eight benchmarking methods to assess their capability in aligning embedding space. Before comparison, we merged the Leiden clusters of each method with the ground truth using a maximum matching strategy for certain categories to produce final clustering results (Supplementary Fig. S1b-f). spatiAlign achieved the highest adjusted Rand index (ARI)^34^ score with a mean of 0.5967 on all four sections and outperformed all the control methods (Fig. 2b). In addition, spatiAlign achieved the highest mean weighted F1 score of the local inverse Simpson’s index (LISI)^8^ of 0.8402 (Fig. 2c), where sufficient mixing and variation preservation were equally evaluated. In comparison, MNN showed ineffectiveness in fusing the sections together and obtained the lowest weighted F1 score of LISI. The uniform manifold approximation and projection (UMAP) visualization for each method revealed that spatiAlign outperformed other control methods in separating clusters while simultaneously integrating slices (Fig. 2d). In particular, methods such as GraphST, SCALEX, Harmony, and Combat did not clearly separate spots belonging to distinct layers, and the batches did not mix well when using MNN. Although PRECAST appeared to separate clusters and integrate batches well, it resulted in Layer_1 being split into two groups.

**Fig. 2.**
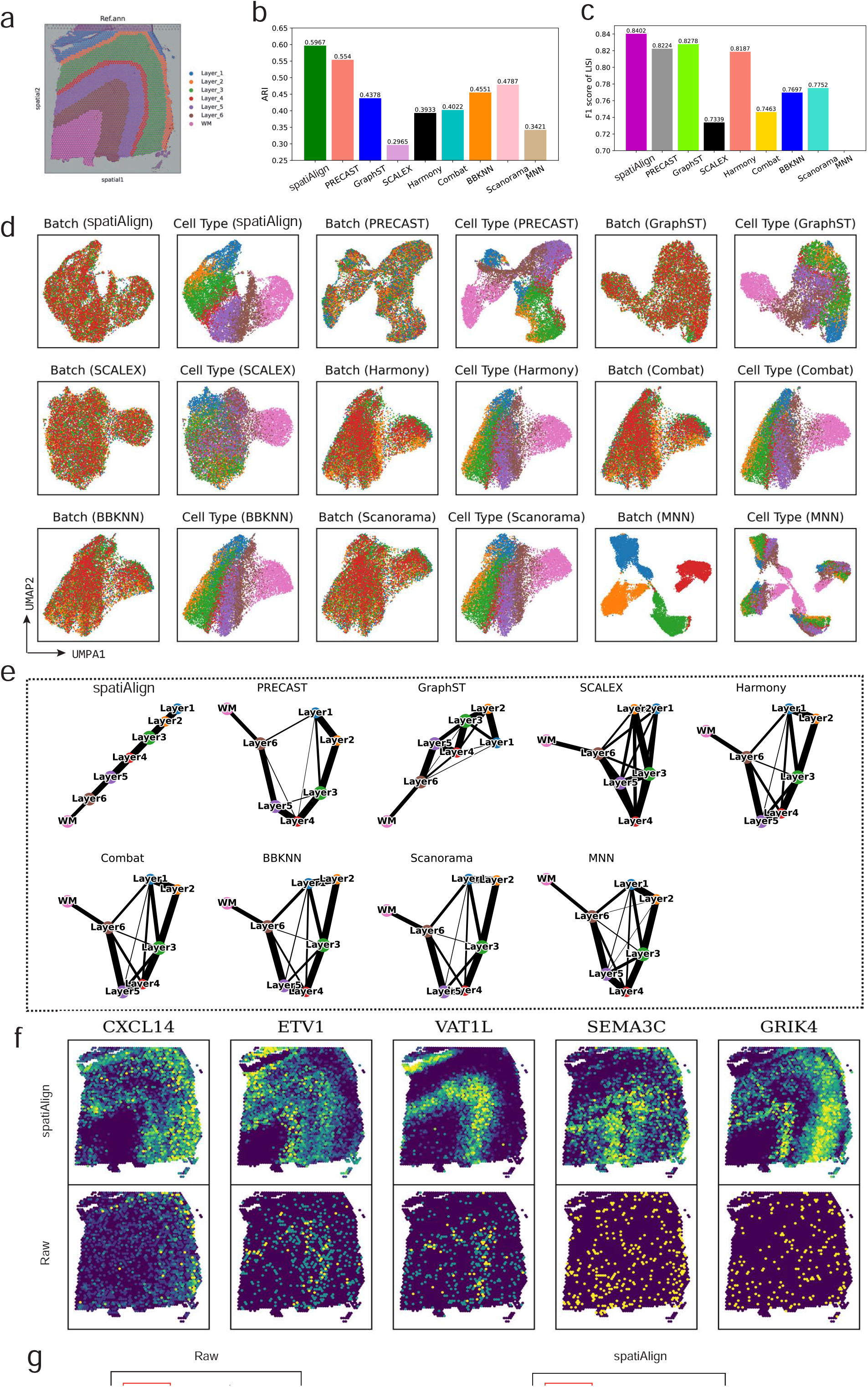
spatiAlign outperforms the control methods in integrating the human dorsolateral prefrontal cortex (DLPFC) datasets. **a)**. Manual annotation of sample ID 151673 from the original study. **b)**. Bar plots of the mean scores of the adjusted Rand index (ARI) for the combined clusters from spatiAlign and other control methods. **c)**. Bar plots of the weighted F1 scores of the local inverse Simpson’s index (LISI), assessing both batch mixing and cell-type separation, for the integration results from different data integration methods. **d)**. UMAP plots for the integrated batches and identified cell types from spatiAlign and other control methods. For the integration result of each method, dots in the right panel are coloured by batch, and dots in the left panel are coloured by cell type. **e)**. PAGA graphs of spatiAlign and other control methods. **f)**. Spatial visualization of spatiAlign-enhanced (top panel) and raw (bottom panel) expression of layer-marker genes. **g)**. Violin plots of the raw (left panel) and spatiAlign-enhanced (right panel) expression of layer-marker genes. The cortical layers corresponding to the layer-marker genes are highlighted with red boxes.

Furthermore, we validated the latent embeddings with the inferred trajectory from PAGA^35^ (Fig. 2e). The PAGA path derived from spatiAlign embeddings exhibited a clear and nearly linear spatial trajectory from Layer_1 to Layer_6, with significant similarities observed between adjacent layers, in accordance with the developmental process of the neurons^36^. In contrast, the PAGA results of the other benchmarking methods were intermixed. Finally, we compared the spatial expression patterns of layer marker genes before and after spatiAlign (Fig. 2f). The results revealed that the spatiAlign-reconstructed expression of layer-marker genes had enhanced laminar enrichment and denoised distributions compared with the original data. For example, *CXCL14* in Layer_1 and Layer_2, *ETV1* in Layer_5 and Layer_6, and *VAT1L* in Layer_5 were consistent with previous studies^37^, whereas their raw gene expression did not show discernible spatial laminar patterns. In addition, violin plots comparing gene expression before and after spatiAlign processing also showed the expression enhancement of spatiAlign (Fig. 2g). For example, the reconstructed expression of *SEMA3C* significantly populated Layer 6 compared with the original data. Such expression enhancements were also observed in other sections, such as in sample ID 151674, further validating the reliability of the reconstructed expressions (Supplementary Fig. S2c).

### spatiAlign enables the alignment of multiple olfactory bulb datasets from different SRT platforms

To demonstrate the efficiency of spatiAlign in integrating datasets from different sequencing platforms, we used three mouse olfactory bulb datasets. One slice was profiled by 10x Genomics Visium, while the other two slices were obtained from Stereo-seq (Fig. 3a). Before integration, we manually annotated each dataset (Fig. 3c) by leveraging unsupervised clustering (Supplementary Fig. S3a, b), reported marker genes (Supplementary Fig. S3c, d, e, f) and the ssDNA image (Fig. 3b). This provided a ground truth for calculating the weighted F1-score of LISI, which quantified the performance of the methods in aligning batches and separating cells from different clusters. As a result, spatiAlign achieved the highest score of 0.7935, outperforming other methods such as PRECAST (0.6863) and SCALEX (0.6099), while MNN was the poorest with a score of 0.0485 (Fig. 3d). Next, on the UMAP plots, we illustrated the batch effects present before alignment (Fig. 3e). After integration, spatiAlign demonstrated successful batch merging, in contrast to the outputs of PRECAST, GraphST, Harmony, Combat and other control methods, where prominent batch effects remained observable. In addition, spatiAlign found separate clusters that aligned well across the three sections (Fig. 3f). Even though BBKNN and SCALEX also generated separate clusters, batch effects were still visible after their integration. Hence, compared with combined clustering results produced by the control methods, those detected using spatiAlign embeddings better corresponded to the annotated ground truth and showed a higher consistency across different sections.

**Fig. 3.**
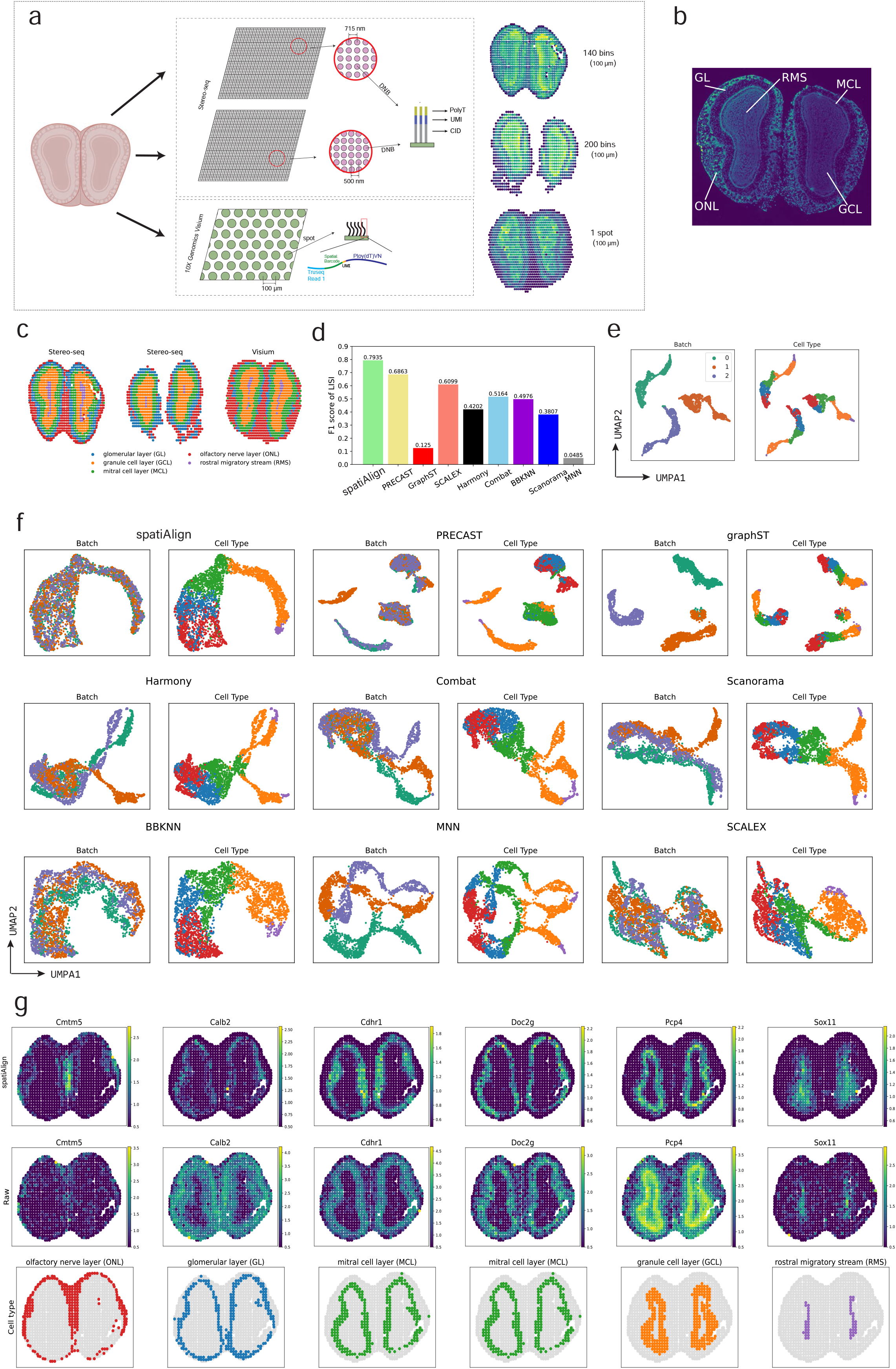
spatiAlign integrates three mouse olfactory bulb datasets from Stereo-seq and 10x Genomics Visium sequencing platforms. **a)**. The mouse olfactory bulb datasets consisted of three sections, with two sections sequenced using Stereo-seq and the third section generated from 10x Genomics Visium. The two Stereo-seq datasets were sequenced on different types of chips, with spots having centre-to-centre distances of 500 and 715 nm, respectively (middle panel). Hence, the two Stereo-seq datasets were individually binned at Bin140 and Bin200 to ensure that all spots in the three sections were of the same size of 100 μm (right panel). **b)**. Organization of mouse olfactory bulb annotated by ssDNA image. **c)**. Manual annotation as a ground truth for benchmarking analysis. Spots are coloured by cell type. **d)**. Bar plots of the weighted F1 scores of LISI for the integration results from spatiAlign and the other control methods. **e)**. Visualization of batch effects present in batches and cell types before integration. **f)**. UMAP plots for the integrated batches and identified cell types from spatiAlign and other control methods. For the integration result of each method, dots in the right panel are coloured by batch, and dots in the left panel are coloured by cell type. **g)**. Spatial visualization of spatiAlign-enhanced (top panel) and raw (middle panel) expression of marker genes, together with the associated cell types (bottom panel). spatiAlign denoised and enhanced the spatial expression pattern of marker genes compared with raw data.

Furthermore, we showed that the reconstructed gene expression from spatiAlign (Fig. 3g, Supplementary Fig. S4d, e, top panel) was denoised and enhanced compared with the raw gene expression (Fig. 3g, Supplementary Fig. S4d, e, middle panel). For some marker genes^38^, e.g., *Cmtm5, Cdhr1, Doc2g*, and *Pcp4*, the spatial expression pattern was clearly enhanced and more consistent with the spatial locations of the corresponding cell types(Fig. 3g, Supplementary Fig. S4d, e, bottom panel).

### spatiAlign preserves heterogeneous characteristics among slices while aligning datasets

We utilized three mouse hippocampal slices from Slide-seq (Fig. 4a and Supplementary Table 1) to assess the performance of spatiAlign and the benchmarking methods in integrating datasets with different biological characteristics. These mouse hippocampus slices were collected from different regions in the mouse brain^3, 39, 40^. As shown on the UMAP plots, spatiAlign accurately integrated disparate datasets and revealed diverse clusters of structural heterogeneity (Fig. 4b, d, and g). Quantitatively, spatiAlign excelled over other control methods with an integrated LISI (iLISI) index of 0.6230, except for SCALEX. However, despite achieving the highest iLISI index, SCALEX was unable to preserve the biological difference among slices (Fig. 4c, Supplementary Fig. S5a and d).

**Fig. 4.**
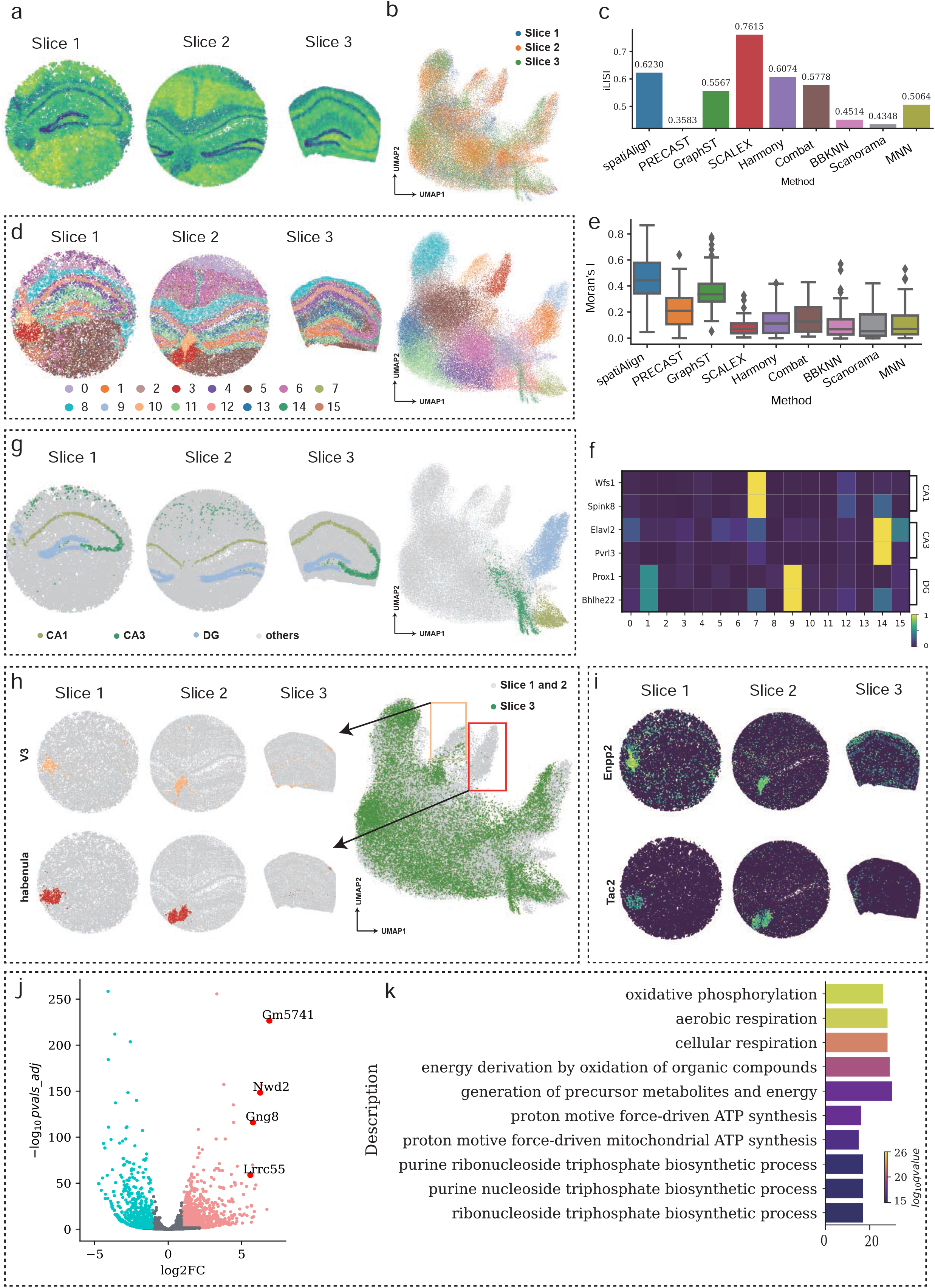
spatiAlign identifies distinct brain structures specific to each slice while integrating three mouse hippocampus datasets. **a)**. Spatial heatmap of total transcripts in the three mouse hippocampal slices measured by Slide-seq. **b)**. UMAP plot for the integrated slices from spatiAlign. **c)**. Bar plots of the integration LISI (iLISI) scores, evaluating batch mixing, for the integration results from spatiAlign and other control methods. **d)**. Spatial visualization (left) and UMAP plot (right) for the joint clustering results from spatiAlign. **e)**. Boxplots of global Moran’s I index for the joint clusters from spatiAlign and other control methods. **f)**. The expression matrix plot of markers of the CA1, CA3, and DG regions. **g)**. Spatial visualization (left) and UMAP plot (right) of CA1, CA3, and DG regions that were only identified by spatiAlign. **h)**. Spatial visualization (left) and UMAP plot (right) of V3 and the habenula that are specific to slice 1 and slice 2. **i)**. Spatial expression of the marker genes *Enpp2* in V3 and *Tac2* in the habenula. **j)**. Volcano plot of differentially expressed genes (DEGs) between the habenula and rest. **k)**. Top ten highly enriched GO terms for the top 100 ranked DEGs.

Furthermore, we adopted hierarchical clustering to validate the effectiveness of each method in identifying the brain regions. The resulting cell clusters after spatiAlign was applied displayed strong spatial aggregation with clear boundaries and higher consistency with the anatomical structures of the Allen Brain Atlas^41^ (Fig. 4d and Supplementary Fig. S5b). Such an observation was further evidenced by the global Moran’s I index, which measures spatial autocorrelation (Fig. 4e). Regarding finding the substructural regions, our proposed spatiAlign was the only method that identified the substructures of the hippocampus, including CA1, CA2 and dentate gyrus (DG), on all three slices (Fig. 4g). The successful hippocampus-related-region identification of spatiAlign had higher consistency across three slices than others (Fig. 4g), while GraphST detected incorrect regions due to a lack of registering spatial coordinates (Supplementary Fig. S5d). For preserving heterogeneous characteristics, we observed that the identified habenula and third ventricle (V3) regions were barely enriched on slice 3 but highly populated on the other two slices, as expected (Fig. 4h). Such results were in high concordance with the expression spatial pattern of the associated marker genes^42^ *Enpp2* for V3 and *Tac2* for habenula (Fig. 4i). To validate the biological traits of heterogeneous embedding, we implemented DEG and GO analyses on detected habenular cell groups. We found many marker genes^42^ for habenula among the highly expressed genes of the merged dataset, e.g., *Gm5741, Nwd2, Gng8* and *Lrrc55* (Fig. 4j). In addition, the GO enrichment analysis showed that the habenula is actively involved in the production and synthesis of ATP (Fig. 4k). This finding was in accordance with biological understandings that ATP not only plays a crucial role in energy metabolism for habenular cells but also acts as a neurotransmitter to modulate neuronal activity and synaptic transmission^43^.

### spatiAlign facilitates joint gene-level analysis of time-series mouse embryonic brain

Finally, we utilized a series of mouse brain datasets^4^ extracted from multiple developing mouse embryos (Fig. 5a), measured by Stereo-seq, to demonstrate the benefits of spatiAlign for downstream gene-level analysis. These brain sections were collected at different embryonic days from E9.5 to E16.5, which included a total of 104,974 cells and 22,864 genes in the merged dataset. Herein, we initially evaluated the inherent batch effects present prior to alignment. Before applying spatiAlign, cells were primarily grouped by batch (Fig. 5b). In comparison, spatiAlign well aligned these datasets within its lower-dimensional representations, where the batch effects were adjusted. The cells were then clustered into coherent groups in an unsupervised manner, and we next manually labelled them by referring to the expression of marker genes reported by the atlas of the developing mouse brain^44^ (Fig. 5b). These marker genes, e.g., *Ccnd2* of NeuB, *Col4a1* of fibroblast, *Sncg* of FMN, *Slc1a3* of Hb VZ, and *Hcrtr2* of Spall VZ, exhibited the highest expression levels in their corresponding cell types that had a relatively high fraction (Fig. 5c). In particular, we found two subtypes of GABAergic interneurons in the subpallial region that were characterized by the *Dlx5* and *Gpm6a genes*, which we named SPall Gpm6a and SPall Dlx5, respectively (Fig. 5c). The validity of these annotations was also confirmed by the strong correspondence observed in the spatial distributions between cell types and relevant marker genes (Supplementary Fig. S6a).

**Fig. 5.**
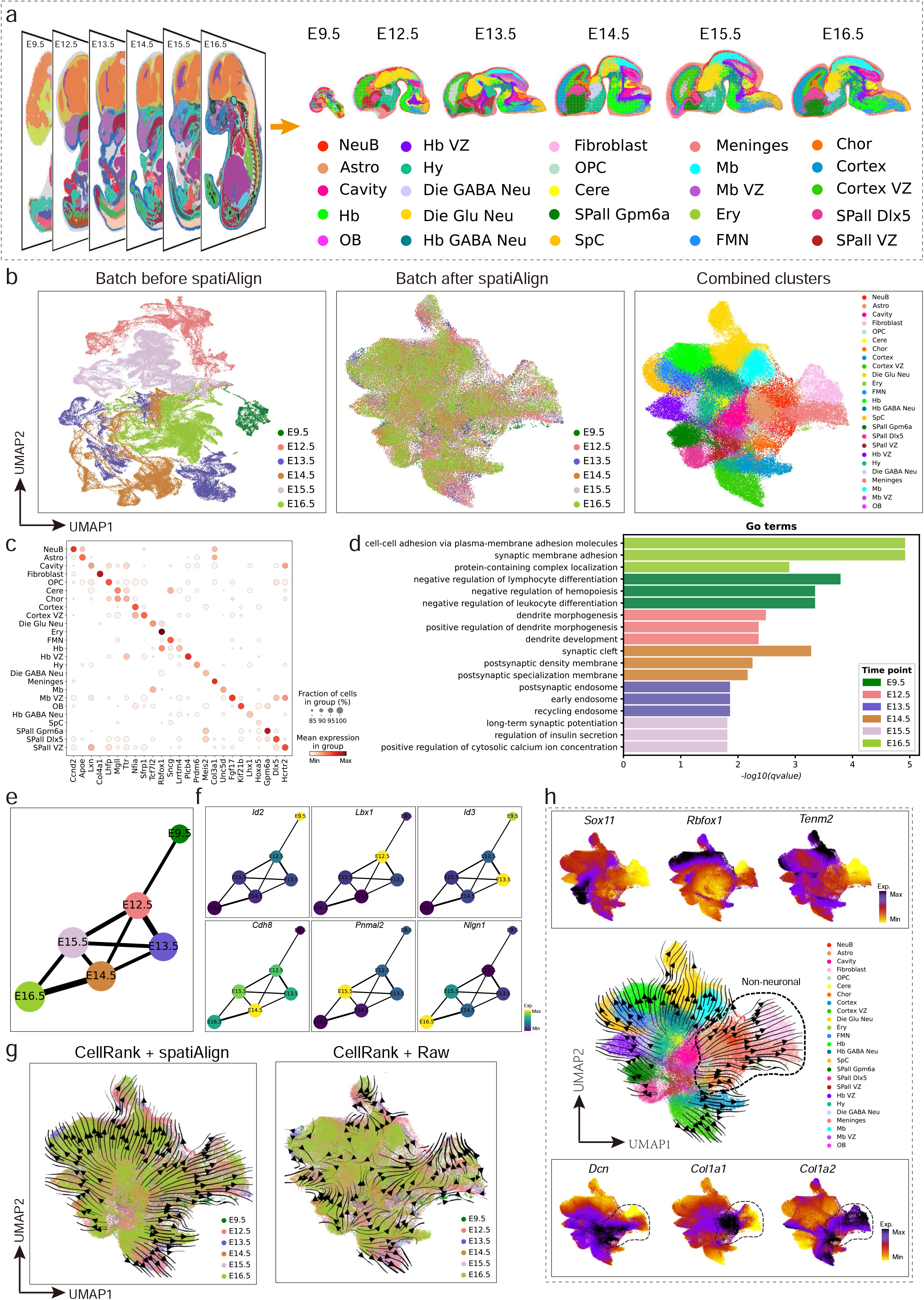
spatiAlign facilitates joint analysis of time-series mouse embryonic brain sections. **a)**. Unsupervised clustering of time-series brain sections extracted from the mouse embryos across E9.5-E16.5 (E9.5, E12.5, E13.5, E14.5, E15.5 and E16.5) after integration using spatiAlign. Spots are coloured by their annotation (right panel). NeuB, neuroblast; Astro, astrocyte; Hb, hindbrain; OB, olfactory bulb; VZ, ventricular zone; Hy, hypothalamus; Die, diencephalon; OPC, oligodendrocyte precursor cell; Cere, cerebellum; SPall, subpallium; SpC, spinal cord; Mb, dorsal midbrain; Ery, erythrocyte; FMN, facial motor nucleus; and Chor, choroid plexus. **b)**. UMAP plots for batch mixing before spatiAlign (left) and after spatiAlign (middle) and the labelled combined clusters from spatiAlign (right). **c)**. Expression dot plots showing the gene expression specificity of typical marker genes for identified cell types. Dot size represents the proportion of expressing cells, and colour indicates the average expression level in each identified cell type. **d)**. Top three highly enriched GO terms for differentially expressed genes from E9.5 to E16.5. **e)**. PAGA graph of spatiAlign embeddings. Each node represents a batch that is connected by weighted edges that quantify the connectivity between batches. **f)**. Age-specific genes traced along the PAGA graph paths. **g)**. Cellular trajectory across different time points inferred by the spatiAlign-corrected feature matrix (left) and raw expression (right), with black arrows representing transition trends. **h)**. Cellular state transitions across cell types (middle panel) and expression of reported driver genes for neuronal (top panel) and nonneuronal cells (bottom panel).

A key benefit of our proposed spatiAlign is its ability to obtain aligned gene expression with batch effects removed, thereby enabling downstream gene-level analysis. Based on the reconstructed expression features, we identified DEGs across E9.5-E16.5 using the Wilcoxon test in SCANPY. A heatmap of the expression of the top 5 ranked DEGs (Supplementary Fig. S6b) illustrated high specificity across different developmental stages. In our observations, the detected DEGs, e.g., *Id2, Lbx1, Id3, Cdh8*, and *Nlgn1*, have been reported to play crucial roles in neuronal differentiation and maturation processes, such as neurogenesis and synaptic plasticity. Specifically, *Id2*, with differential expression at E9.5, has been extensively studied for its involvement in balancing neuronal proliferation and differentiation^45^. Similarly, *Id3*, showing specificity to E13.5, was widely recognized for its function in controlling the timing of neurogenesis in the embryo^46^. Conversely, the top-ranked DEGs identified at E16.5, such as *Nlgn1, Cadm2, Nrg1*, and *Ccser1*, have been well studied for their contributions to synapse formation, myelination, synaptic plasticity and connectivity^47-49^, suggesting the final stage of neurogenesis with synaptogenesis and the formation of synaptic connections between neurons at E16.5. The subsequent GO-based enrichment analysis (Fig. 5d) revealed distinct functional enrichments during different developmental stages in the mouse embryonic brain. Negative regulation of haemopoiesis was observed at E9.5, followed by dendrite morphogenesis at E12.5, early endosome at E13.5, synaptic cleft at E14.5, long-term synaptic potentiation at E15.5, and synaptic membrane adhesion at E16.5. These findings were in line with the major developmental events observed at different embryonic stages, encompassing the initiation of neurogenesis (E9.5)^50^, early neuronal connection (E12.5)^51^, increased neurogenesis (E13.5, E14.5 and E15.5)^52, 53^, and the refinement of synaptic plasticity (E16.5).

We further demonstrated the effectiveness of spatiAlign for combined trajectory analysis by employing two distinct approaches: PAGA, a graph abstraction technique based on low-dimensional embedding space^35^, and CellRank^54^, a state-of-the-art cell fate mapping algorithm using a high-dimensional count matrix as input. The PAGA graph of spatiAlign embeddings (Fig. 5e) exhibited a nearly linear development trajectory from E9.5 to E16.5, as well as a high similarity between adjacent time points. Furthermore, the age-specific genes that were identified could be traced along the PAGA path (Fig. 5f). We proceeded to compare the reconstructed trajectory obtained from CellRank using two different inputs: the raw and spatiAlign-reconstructed feature matrices. The recovered trajectory, derived from reconstructed features (Fig. 5g and h), illustrated a clear transition path across cell types and a similar distribution across different time points, consistent with previous observations^55^. In contrast, the batch effects present in the raw count matrix may lead to infeasible and chaotic fate potentials across different batches (Supplementary Fig. S6c). Additionally, the expression patterns of reported driver genes associated with neuronal (i.e., *Tuba1a*^56^, *Tenm2*^57^, *Rbfox1*) as well as nonneuronal (*Dcn, Col1a1, Col1a2*) development^58^ (Fig. 5h) were consistent with the predicted cell fate, thereby validating the feasibility of the estimated pseudotime and affirming the reliability of our analysis.

## Discussion

In this paper, we develop spatiAlign, an advanced deep learning methodology that tackles the challenge of integrating multiple SRT datasets. SpatiAlign first transforms spatial information into a neighbouring adjacency matrix to perform spatial embedding that aggregates gene expression profiles together with spatial neighbouring context for spot/cell representations. The obtained representations are subsequently fine-tuned through augmentation-based contrastive learning, which incorporates spatial context information to improve their informativeness and distinguishability. Next, regarding aligning biological effects, spatiAlign adopts across-domain adaptation and deep clustering strategies to bring the semantic similarity of spots/cells closer and push dissimilar spots/cells apart, regardless of which datasets they are from. Collectively, beyond SRT dataset integration and batch effect correction, spatiAlign-integrated datasets can be used for downstream analysis, such as identifying combined clusters and DEGs and trajectory inference.

Naturally, one might be concerned that achieving a sufficient mix of serial tissue sections could result in the inability to distinguish spots/cells from different clusters. Therefore, in this study, we introduce a weighted F1 score of LISI, which evaluates the integration mixing and separation of each cluster, to perform comparison analysis. We presented a series of benchmarking analyses on four publicly available SRT datasets with different characteristics. On the human DLPFC datasets, with the manual annotation as ground truth, spatiAlign achieves the highest ARI and weighted F1 score of LISI compared with other control methods. This quantitative assessment highlights its superiority in integrating different samples while also identifying separate clusters. Furthermore, the superior performance of spatiAlign on aggregated datasets of olfactory bulbs sequenced by different platforms demonstrates its efficacy in integrating multiple datasets with complex technical variations. In addition, the reconstructed expression of region-specific marker genes exhibits a greater spatial specificity compared with the original data. However, we point out here that our effort was not intended to develop a new imputation method over existing methods but to demonstrate that spatiAlign-reconstructed matrices enhance gene counts.

Moreover, there is concern regarding the potential loss of distinct biological characteristics during the batch alignment process. Herein, we unequivocally affirm that spatiAlign not only effectively preserves the intrinsic variation among sections but also adeptly harmonizes batches, as demonstrated through its successful application to three distinct brain sections characterized by heterogeneous structures. However, the benchmarking methods are unable to match the performance of spatiAlign. When applied to a time-series dataset, spatiAlign significantly facilitates downstream analysis, such as combined clustering, combined differential expression analysis and trajectory inference. In the results, various subtypes of neurons were successfully identified, with the typical marker genes displaying the highest expression in their corresponding cell types. Upon analysing the reconstructed full expression space, we identified DEGs and significant GO terms specific to different developmental stages that showed high consistency with previous studies on mouse brain development. Comparing the trajectories inferred from corrected expression features and the raw data, we verify that spatiAlign not only aligns multiple batches into a joint low-dimensional embedding space but also corrects the batch effects in their full expression space. This capability empowers users to perform preprocessing for methods that require a full gene expression matrix, such as CellRank.

We designed spatiAlign to be user-friendly and believe that it offers a novel and effective approach for SRT dataset integration. In the future, we envision extending spatiAlign for integrative and multimodal spatial molecular dataset analysis, e.g., epigenetics, proteomics and microbiomics. Such advancements will enable efficient integration of multiomics data and facilitate the deeper exploration of biological phenomena.

## Methods

### Motivation for the use of across-domain adaptation contrastive learning

As genomic sequencing technology continues to advance, an increasing number of SRT datasets are being generated from various platforms. Joint analysis of multiple datasets can be used to facilitate the extraction of maximum reliable information, but inconsistent data distributions between different sections due to batch effects may affect the reliability of downstream analysis results. To address this issue and maximize the preservation of biological variations, it is desirable to amalgamate disparate datasets and bring similar cell types closer together while keeping dissimilar cell types far apart. Across-domain adaptation contrastive learning, an unsupervised domain adaptation method, can be used for this purpose. This method can align data distributions, preserve biological variations, and remove batch effects while also incorporating spatial information of the SRT dataset into the newly generated latent embedding and reconstructed matrix.

### Data preprocessing

spatiAlign utilizes a series of gene expression matrices and associated spatial coordinates as inputs. The gene expression profiles are stored in a *X*^*N*×*D*^ matrix of unique molecular identifier (UMI) counts, where *N* is the number of spots/cells and *D* is the number of genes, and it also includes (*x, y*) two-dimensional spatial coordinates for each spot/cell. The raw gene expression matrices were first filtered according to criteria *min*_*genes =* 20 and *min*_*cells =* 20 for each dataset using SCANPY (version: 1.9.1), followed by normalization and log transformation of individual spots.

### Spatial neighbour graph construction for the SRT dataset

To fully exploit the spatial local neighbouring context, we convert the spatial coordinates into an undirected neighbourhood graph *G* (*V,E*) by Euclidean distance with a predefined neighbour parameter *k*, where *V* represents the SRT dataset spots/cells and *E* represents the connected edges between the current spot/cell and neighbouring spots/cells. The adjacency matrix of graph *G* is denoted by *A*, in which spot/cell *u*∈*V* with *k* nearest neighbour spots/cells; if spot/cell *v*∈*V* is the neighbour of spot/cell ***u***, *a*_*uv*_ = 1 ; otherwise, it is 0. Specifically, we selected the top 15 nearest neighbours for each spot/cell in the SRT gene expression spatial coordinates.

### Batch-specific variations to separate using domain-specific batch normalization

Batch normalization (BN)^59^ is widely used to solve the problem of internal covariate shift during DNN training. It can reduce the problems of vanishing gradients and overfitting. For a mini-batch of data ℬ *= x*_1…m_, the BN layer can be calculated using the following parameterization:

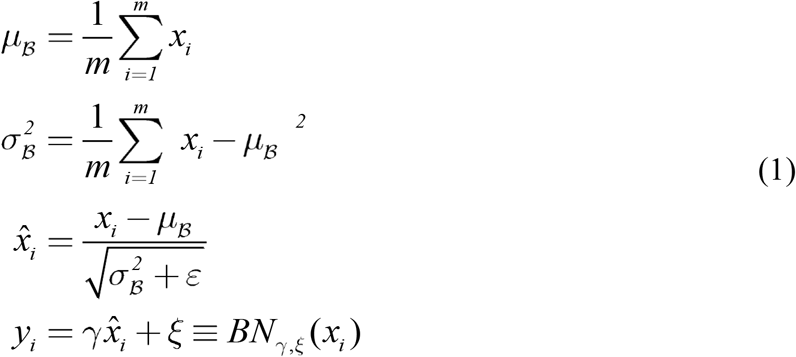

where *μ*_ℬ_ is the mean of the mini-batch, 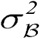 is the variance of the mini-batch, 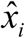 is the normalized output by the mean and variance of the mini-batch, *ε* is a small constant to avoid dividing by zero, and *y*_*i*_ is the output of the BN layer, which is obtained by scaling and shifting 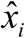 with learned parameters *γ* and *ξ*.

Domain-specific batch normalization (DSBN)^60^ is used in unsupervised domain adaptation with multiple source datasets to separate domain-specific variations from different datasets. In spatiAlign, DSBN consists of multiple sets of BN layers that select the corresponding BN with the batch label *b*. DSBN can be represented as follows:

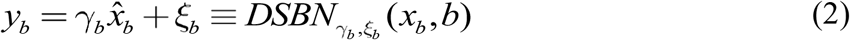

where *γ*_*b*_ and *ξ*_*b*_ are batch-specific affine parameters for batch *b*.

DSBN has been proposed to capture and utilize the batch-specific information in datasets by learning affine parameters for each dataset, which enables the model to learn the batch-specific variations that exist within the datasets^13, 60^.

### DGI-based feature extractor for reducing dimensions and propagating spatial neighbouring context

spatiAlign leverages the DGI framework to project a gene expression matrix into a latent space while simultaneously performing dimension reduction and propagating spatial neighbouring spots/cells context. To reduce the dimension of higher-dimensional SRT data, we employed a DNN-based autoencoder mapping model *f*_*θ*_ : *X* → *Z*, where *θ* represents the parameters of the mapping model, *Z*^*N*×*d*^ is a latent embedding with dimensions *d*, and *d* ≪ *D*. The DNN-based mapping model, a feature embedding block, consists of a fully connected block and two stacked residual bottleneck blocks. Specifically, the fully connected block comprises a linear connected layer, a DSBN layer, an exponential linear unit (ELU) as a nonlinear activation function, and a dropout layer in sequence. Each residual bottleneck block consists of two stacked fully connected blocks, and the output of the residual bottleneck block is passed through an ELU layer (Fig. 1b). Notably, the feature embedding block only takes the gene expression matrix as input.

To propagate the spatial neighbouring context in the reduced dimensionality space, we employ a variational graph autoencoder (VGAE) framework. The VGAE framework takes the latent embedding *Z* obtained from the feature embedding model and the adjacency matrix *A* as input and generates *Y* as output. The VGAE encoder includes two stacked graph convolutional network (GCN) layers and uses the rectified linear unit (ReLU) as a nonlinear activation function. The first GCN layer generates a lower-dimensional spatial embedding and aggregates the spatial neighbouring context, while the second GCN layer generates the mean *μ* and variance *δ*^*2*^. The spatial embedding *Y* is then reparametrized from *Y =μ* + *τ* ∗ *δ*^*2*^, where *τ* ∼ *N* (0,1). The final latent representation *S* is generated from the feature fusion block, which includes two stacked fully connected layers, as well as a DSBN layer followed by each connected layer in sequence, and takes concatenated feature embedding as input, which is obtained by concatenating the reduction dimensionality embedding *Z* and the spatial embedding *Y*. The final latent embedding *S* is then used to reconstruct the original gene expression matrix *X*′ in the DNN-based autoencoder and the spatial neighbouring adjacency matrix *A*′ in the VGAE network.

Training the DNN-based autoencoder and VGAE network minimizes the loss of the reconstructed gene expression matrix and maximizes the log-likelihood of the observed SRT sequencing latent representation *S*. We first employed the scale-invariant mean squared error (MSE)^61^ to measure the DNN-based loss. In addition, the loss function of the VGAE includes a binary cross-entropy loss to minimize the difference between the input spatial neighbouring adjacency matrix *A* and the reconstructed adjacency matrix *A*′. Additionally, a Kullback‒ Leibler divergence loss was used to optimize the log-likelihood between the posterior distribution *q*_*θ*_ (*Y* | *S, A*) and prior distribution *p*(*Y*), where *p*(*Y*) ∼ *N* (0,1). The dimension reduction and spatial neighbouring context propagation loss can be calculated as follows:

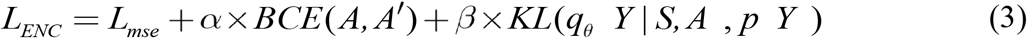

where *BCE* is the binary cross-entropy, *KL* is the Kullback‒Leibler divergence, *L*_*sim*_ *mse*_ is the scale-invariant MSE and *α, β*∈ 0,1 are hyperparameters.

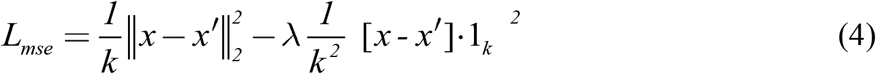

where *k* is the number of spots/cells in the input gene expression matrix, 1_*k*_ is a vector of ones of length *k*, 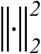 is the squared *L*_*2*_ norm, and *λ* ∈ 0,1 is a hyperparameter.

### Self-supervised contrastive learning for representation enhancement

DGI is a self-supervised learning architecture that maximizes mutual information between local neighbours of a graph to learn representations of nodes. spatiAlign takes original and corrupted gene expression matrices as inputs and generates latent representation matrices *S* and *S*′, respectively. The corrupted matrix is a rowwise random perturbation of the original matrix, and we assume that the corrupted gene expression profiles have the same neighbouring adjacency matrix as the original profiles. Formally, given a spot *i*, we form a positive pair consisting of its representation *s*_*i*_ and the neighbouring graph spot vector *g*, while the corresponding corrupted representation 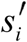 from the corrupted matrix and the same neighbouring graph spot vector *g* form a negative pair. A self-supervised contrastive learning method was used to train the DGI framework, and the loss function was designed to maximize the mutual information of positive pairs while minimizing the mutual information of negative pairs:

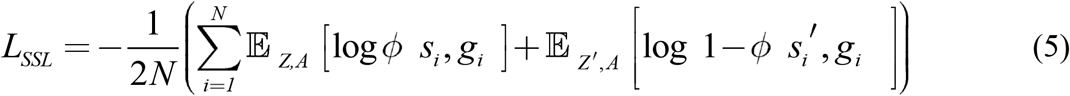

where *ϕ* · is a discriminator, a bilinear layer and follows a sigmoid layer, to distinguish the positive pairs from negative pairs.

### Biological effects alignment using across-domain adaptation contrastive learning

To align biological effects using across-domain adaptation contrastive learning, we propose a criterion for forming pairs based on the assumption that datasets from multiple tissue sections share at least one common cell type in the current alignment setting. To achieve this, we perform in-batch instance-level contrastive learning and across-batch instance-level contrastive learning for each tissue section separately. Specifically, we maintain a memory bank *V*^*b*^ for each tissue section, which is used to store the latent embedding and prototype spot/cell type representations within the batch.

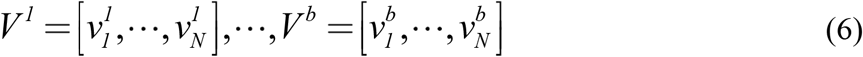

where *v*_*i*_ is the stored feature vector of *x*_*i*_, initialized with final latent representation *S*, and updated with a momentum *m* after each iteration for each dataset:

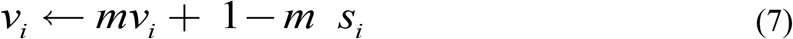

#### In-batch instance level contrastive learning

The pairwise similarity distributions 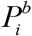 are measured by the cosine distance between latent embedding *S*^*b*^ and the corresponding memory bank *V*^*b*^ to perform in-batch instance discrimination,

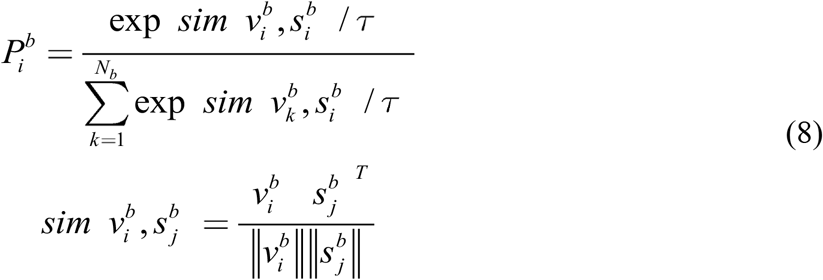

where *τ* is the temperature parameter, which can determine the concentration level of the similarity distribution. Finally, cross-entropy was employed to minimize the in-batch instance discrimination.

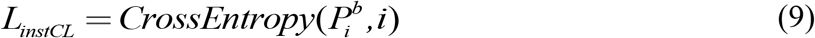

where *i* denotes the unique index of the spot of *x*_*i*_.

#### Pseudoprototypical cluster level contrastive learning

Inspired by unsupervised contrastive clustering^32^, we map each spot/cell *i* into an embedding space with *d* dimensions, where *d* is equal to the number of pseudoprototypical clusters. Since each spot belongs to only one cluster, ideally, the row of the latent embedding *S*^*N*×*d*^ tends to be one-hot, meaning that the *j*-th column of *S*^*N*×*d*^ represents the *j*-th cluster. Similar to in-batch instance-level contrastive learning, our method uses cosine distance to measure the similarity between latent embedding and the corresponding memory bank and maximize the pseudo cluster pair similarity using cross-entropy. Specifically, the loss function can be expressed as:

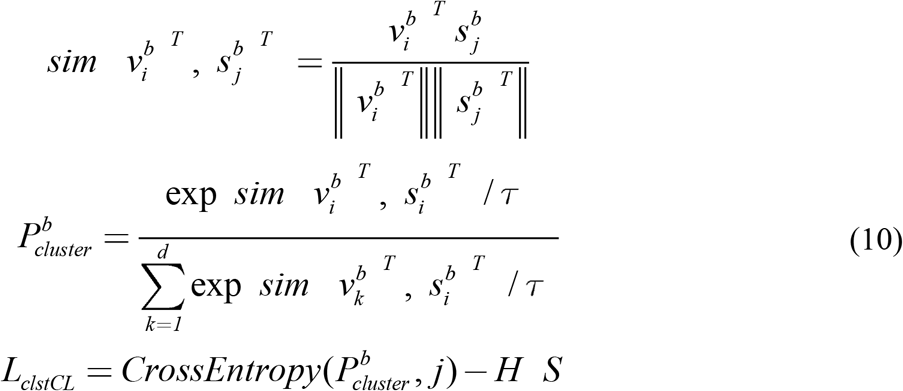

where 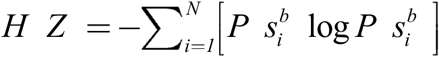 is the entropy of the pseudo cluster assignment probabilities 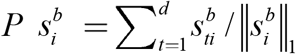, which can help to avoid the trivial solution in which most spots are assigned to the same cluster^32^.

#### Across-batch instance self-supervised learning

To explicitly align biological effects and ensure that spatiAlign learns discriminative representations of dissimilar cell types between different batches, we perform across-batch feature matching. Specifically, we minimize the entropy of the pairwise similarity distribution between latent embeddings in one batch and the latent embeddings stored in the memory bank of another batch. The loss function for across-batch spot/cell pair matching can be formalized as:

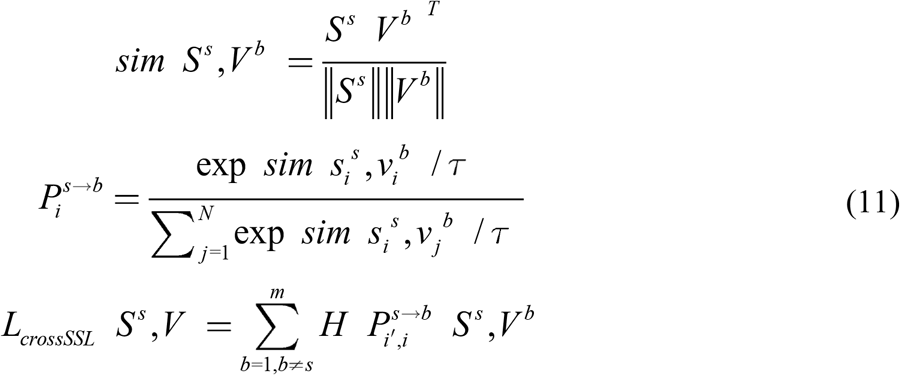

The overall objective for spatiAlign is to minimize:

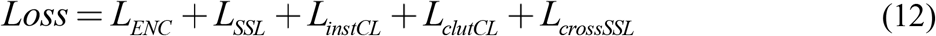

### Comparisons of methods

We perform four comprehensive representative SRT datasets with varying characteristics to compare spatiAlign with other state-of-the-art methods of data integration.

We applied the following integration methods: (1) Harmony^8^ implemented in the SCANPY package external module *harmony_integrate*; (2) Combat^62^ implemented in the SCANPY package module *combat*; (3) Scanorama^11^ implemented in the SCANPY package external module *scanorama_integrate*; (4) BBKNN^12^ implemented in the SCANPY package external module *bbknn*; (5) MNN^15^ implemented in the SCANPY package external module *mnn_correct*; (6) SCALEX^13^ implemented in the Python package *scalex*, and spatial-base methods: (7) PRECAST^24^ implemented in the R package *PRECAST*; (8) GraphST^25^ implemented in the Python package *GraphST*. We input the preprocessed datasets into spatiAlign and several other tested methods. The first six methods were developed for scRNA-Seq datasets, whereas PRECAST and GraphST were specifically designed for SRT datasets.

### Evaluation metrics

We evaluate the performance of spatiAlign and other control methods in both data integration and the preservation of biological variation using the following metric.

#### F1-score of Local inverse Simpson’s index

To simultaneously evaluate the separation of same-cell-type aggregation and across-batch fusion in the data integration, we calculated the LISI^8^ using two different groupings: (1) grouping using different datasets as the batch *iLISI* and (2) grouping using known cell types as the spot *cLISI*. In the data integration, a larger value of *iLISI* indicates sufficient mixing of the different batch datasets, while a smaller value of *cLISI* suggests better preservation of the biological variations between spot types.

The two metrics can be summarized using the *F*1 score as follows:

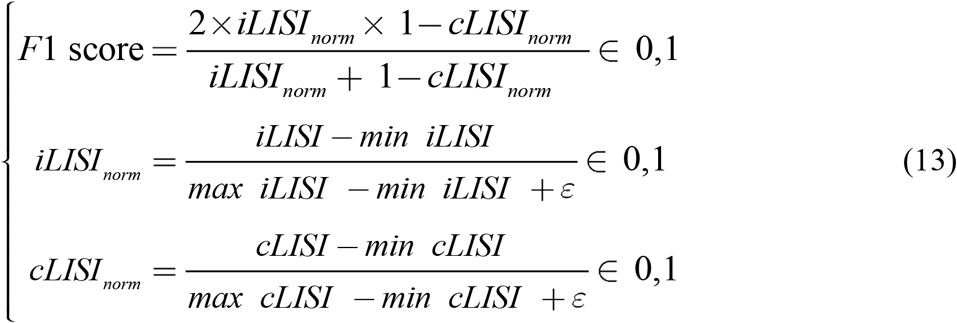

where *ε* is a smaller constant. A higher *F*1 score indicates superior data integration, which effectively retains the biological variations between spot types while eliminating other noncellular biological variations across multiple batches, thereby enhancing the fidelity of the biological information.

#### Adjusted Rand index

To evaluate the efficacy of merge clustering when utilizing lower-dimensional gene expression representations, we utilized the adjusted Rand index (ARI)^34^ as a performance metric. ARI represents an enhanced version of the Rand Index (RI), which overcomes several of its limitations. By measuring the degree of similarity between two partitions, ARI provides a numerical value that ranges between −1 and 1, with a higher value indicating a higher degree of similarity between the two partitions being compared. Moreover, ARI attains a value of 1 when the two partitions under comparison are equivalent up to a permutation. Hence, ARI serves as a reliable and robust tool for evaluating the performance of merge clustering approaches.

#### Hierarchical clustering and Moran’s I index calculation

The spatial regions were identified by a hierarchical clustering algorithm with a lower-dimensional representation from different methods. The *agglomerative clustering* function in the scikit-learn package was implemented with 16 clusters (*n_cluster=*16). Then, we calculate the global *Moran’s I index* for each region on each slice. First, the batch labels were encoded to one-hot vectors, and spatial coordinates were used to calculate spatial neighbours (edge weights=1). Then, the Moran function in the ESDA (2.4.3) Python package was applied to calculate *Moran’s I index*

#### Differential expression analysis and GO enrichment analysis

We employed the *FindMarkers*() function of the Scanpy package to identify differentially expressed genes (DEGs) for the spatial domain using “*T test*” implementation and cutting of the adjusted *p* value at 0.05. To perform GO enrichment analysis for the DEGs, we utilized the *ClusterProfiler* (v4.8.1) R package.

#### Trajectory inference analysis

We used the spatiAlign embedding to infer the PAGA^35^ path by the *scanpy*.*tl*.*paga* function in SCANPY. CellRank^54^ was implemented to estimate pseudotime using the *CytoTraceKernel* algorithm and *compute_transition_matrix* beyond RNA velocity because the spliced and unspliced counts were not available in the mouse embryonic brain datasets. We visualized the directed transition matrix CellRank calculated with the same sort of arrows that are used for RNA velocity. However, there is no RNA velocity in this study.

## Dataset availability

The public datasets are freely available as follows. The Stereo-seq data have been deposited into the CNGB Sequence Archive (CNSA) of the China National GenBank DataBase (CNGBdb) with accession number CNP0001543, the spatiotemporal dataset of the mouse embryonic brain is available at https://db.cngb.org/stomics/mosta, and the 10x Genomics Visium data have been published at https://www.10xgenomics.com/resources/datasets/adult-mouse-olfactory-bulb-1-standard. The LIBD human dorsolateral prefrontal cortex (DLPFC) dataset and mouse breast datasets can be downloaded from https://zenodo.org/record/6925603#.YuM5WXZBwuU. Mouse hippocampus: https://singlecell.broadinstitute.org/single_cell/study/SCP815/highly-sensitive-spatial-transcriptomics-at-near-cellular-resolution-with-slide-seqv2#study-summary, https://singlecell.broadinstitute.org/single_cell/study/SCP354/slide-seq-study#study-summary, and https://singlecell.broadinstitute.org/single_cell/study/SCP948/robust-decomposition-of-cell-type-mixtures-in-spatial-transcriptomics#study-summary, respectively.

## Codes & Software availability

An open-source Python implementation of spatiAlign and reproduction codes are available at: https://github.com/STOmics/Spatialign.git

Tutorials are available at: https://spatialign-tutorials.readthedocs.io/en/latest/index.html

## Acknowledgements

We thank China National GeneBank for providing data support for this study. We thank Guangdong Bigdata Engineering Technology Research Center for Life Sciences support for this study. In addition, we would like to thank Prof. Dr. Junjun Jiang, Dr. Jian Zhang, Dr. Ke Fan, Dr. Yong Bai and Dr. Min Xie for their help.

## Author contributions

Conceptualization: Chao Zhang.

Project administration and supervision: Ao Chen, Xun Xu, Yong Zhang and Yuxiang Li.

Algorithm development and implementation: Chao Zhang.

Public datasets collection, processing and application: Chao Zhang, Lin Liu, Ying Zhang, Mei Li and Shuangsang Fang.

Methods comparisons: Chao Zhang, Lin Liu and Ying Zhang.

Biological interpretation: Chao Zhang, Lin Liu and Ying Zhang.

Manuscript writing and figure generation: Chao Zhang, Lin Liu and Ying Zhang.

Manuscript reviewing: Shuangsang Fang, Qiang Kang and Mei Li

All authors approved the manuscript.

---

**Supplementary Fig. S1.**
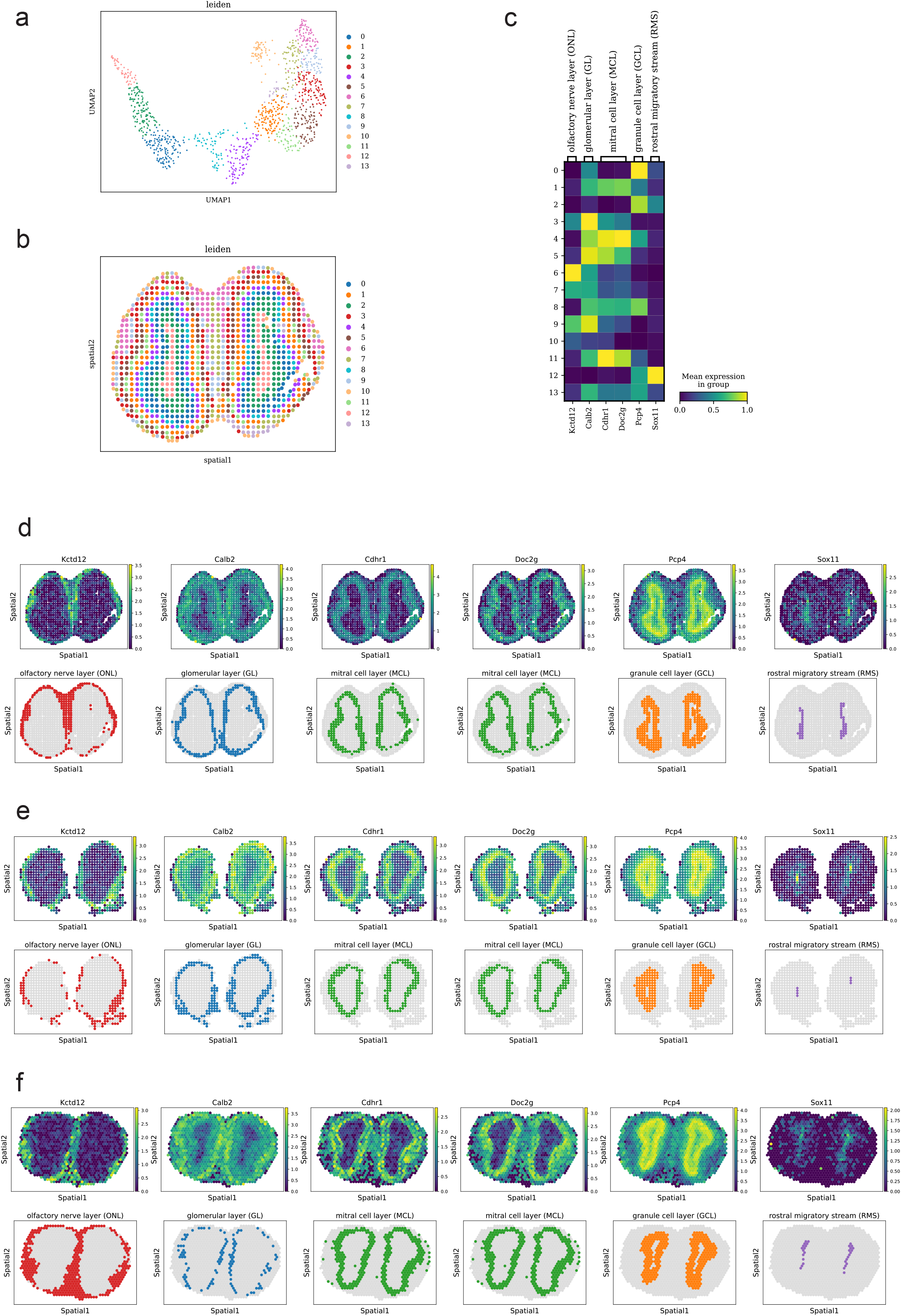
Manual annotation of human DLPFC datasets and joint clustering results from spatiAlign and other control methods, related to Figure 2. **a)**. Manual annotation of four DLPFC sections from the original study. **b)**. UMAP plots for joint leiden clusters (Leiden) from spatiAlign and the control methods, together with the final clusters (Mapping) that merged leiden clusters with the ground truth using a maximum matching strategy. **c, d, e, f)**. Spatial visualization of the Leiden clusters and the mapping clusters of sample ID 151673 (**c**), sample ID 151674 (**d**), sample ID 151675 (**e**), and sample ID 151676 (**f**).

**Supplementary Fig. S2.**
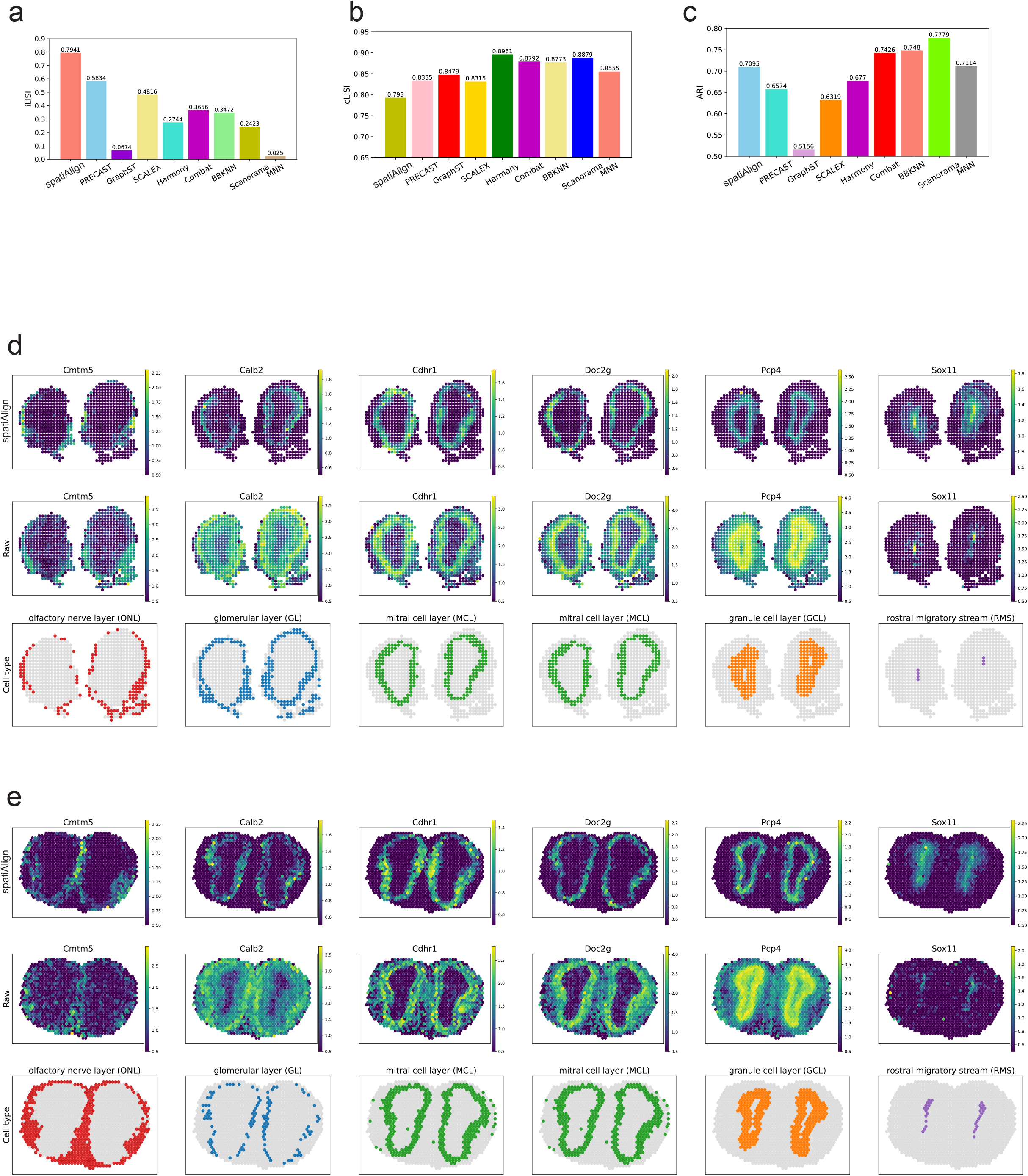
Benchmarking analysis on human DLPFC datasets, related to Figure 2. **a, b)**. Bar plots of integration LISI (iLISI), **a**) and cell-type LISI (cLISI), **b**) scores for integration results from different methods. **c)**. Visualization of spatiAlign-enhanced (top panel) and raw (bottom panel) spatial expression of layer-marker genes in sample 151674.

**Supplementary Fig. S3.**
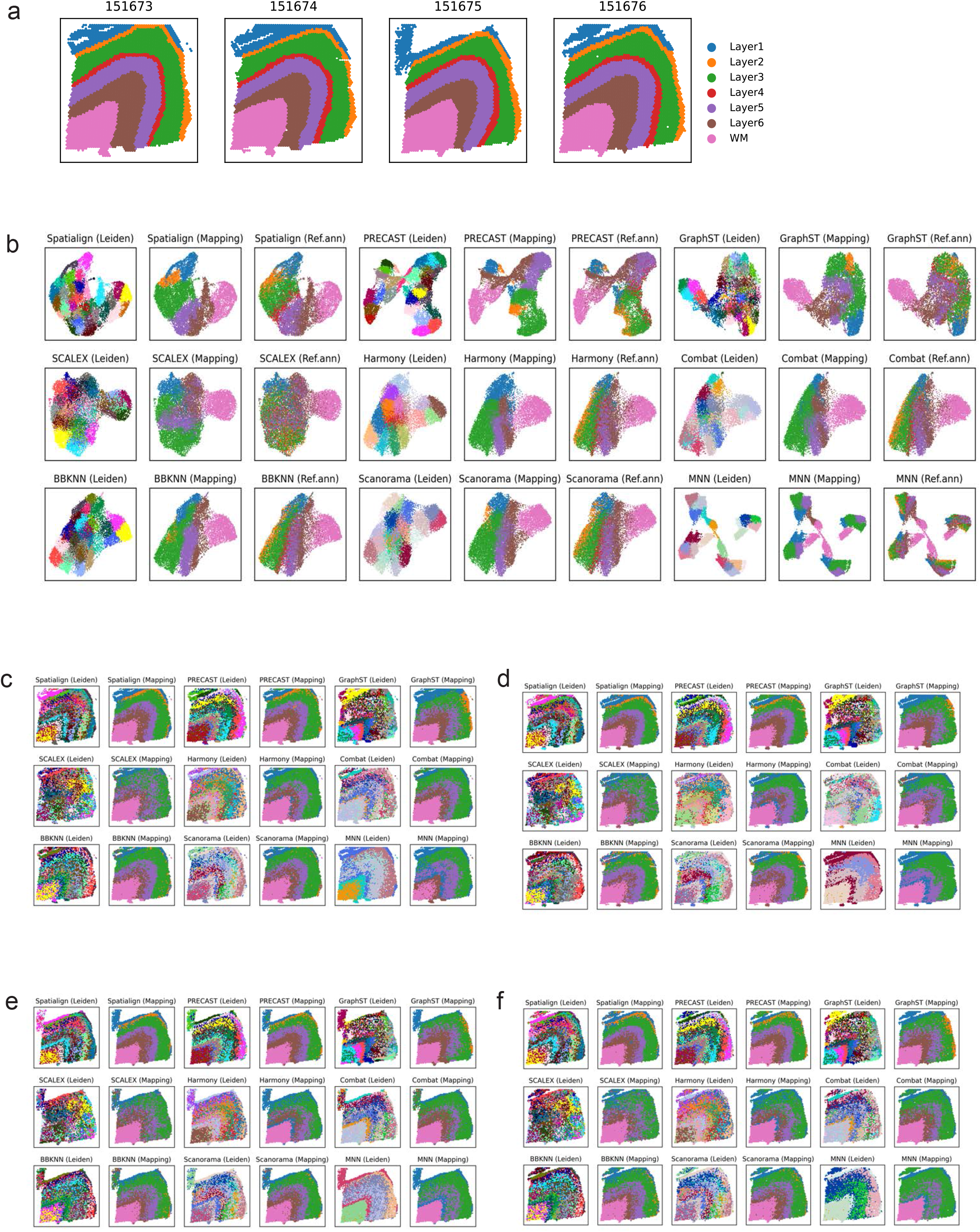
Manual annotation of olfactory bulb datasets, related to Fig. 3. **a)**. UMAP plot for the left clusters of a Stereo-seq olfactory bulb dataset and its spatial visualization (**b**). **c)**. Heatmap of marker genes associated with their cell types. **d, e, f)**. Spatial pattern of marker genes and the corresponding cell types on the three olfactory bulb slices.

**Supplementary Fig. S4.**
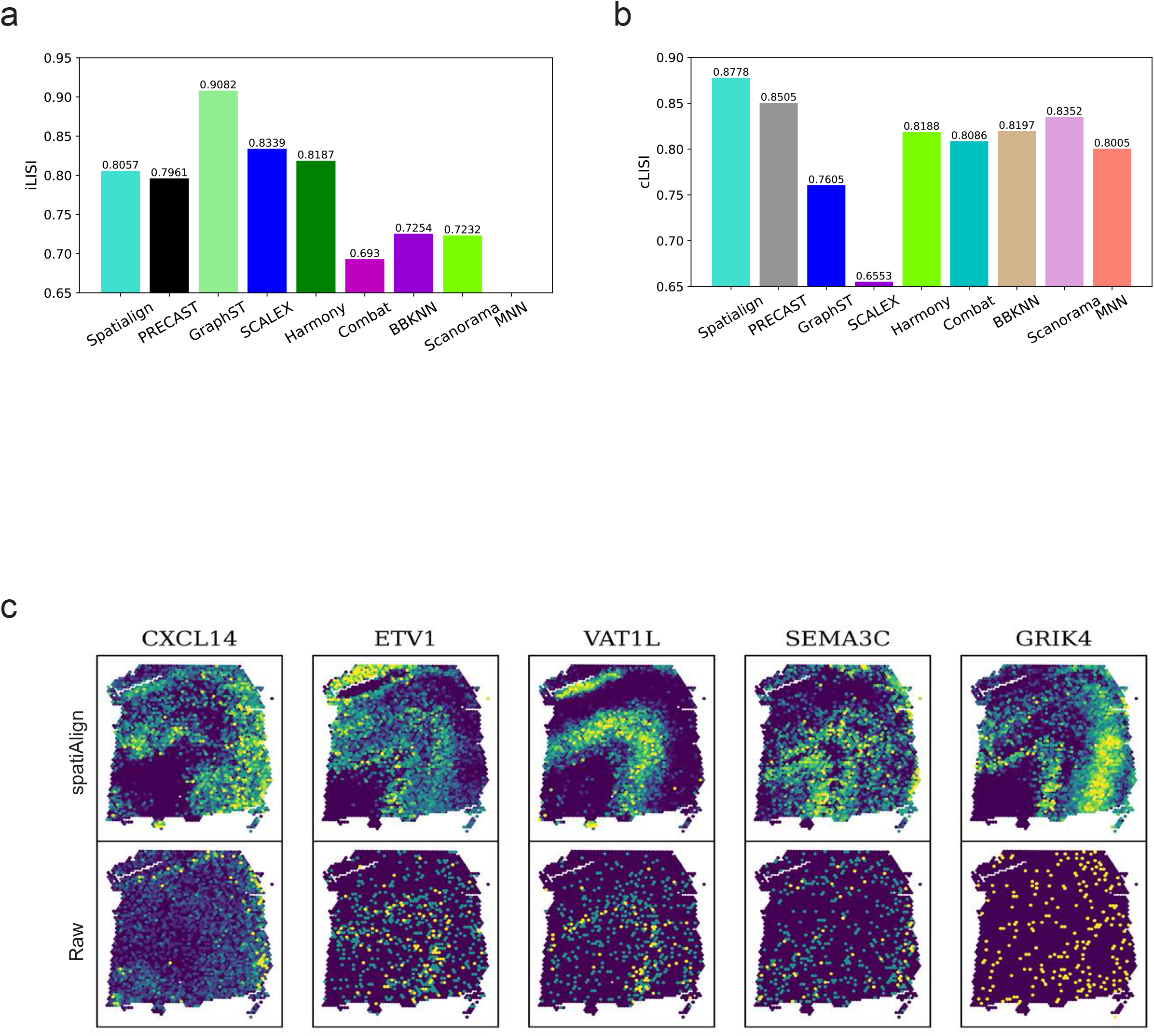
Benchmarking analysis on olfactory bulb datasets, related to Figure 3. **a, b, c)**. Bar plots of integration LISI (iLISI), **a**), cell-type LISI (cLISI), **b**) and ARI (**c**) scores for integration results from different methods. **d, e)**. Spatial visualization of spatiAlign-enhanced (top panel) and raw (middle panel) spatial expression of marker genes, together with their corresponding cell types (bottom panel), on two olfactory bulb sections.

**Supplementary Fig. S5.**
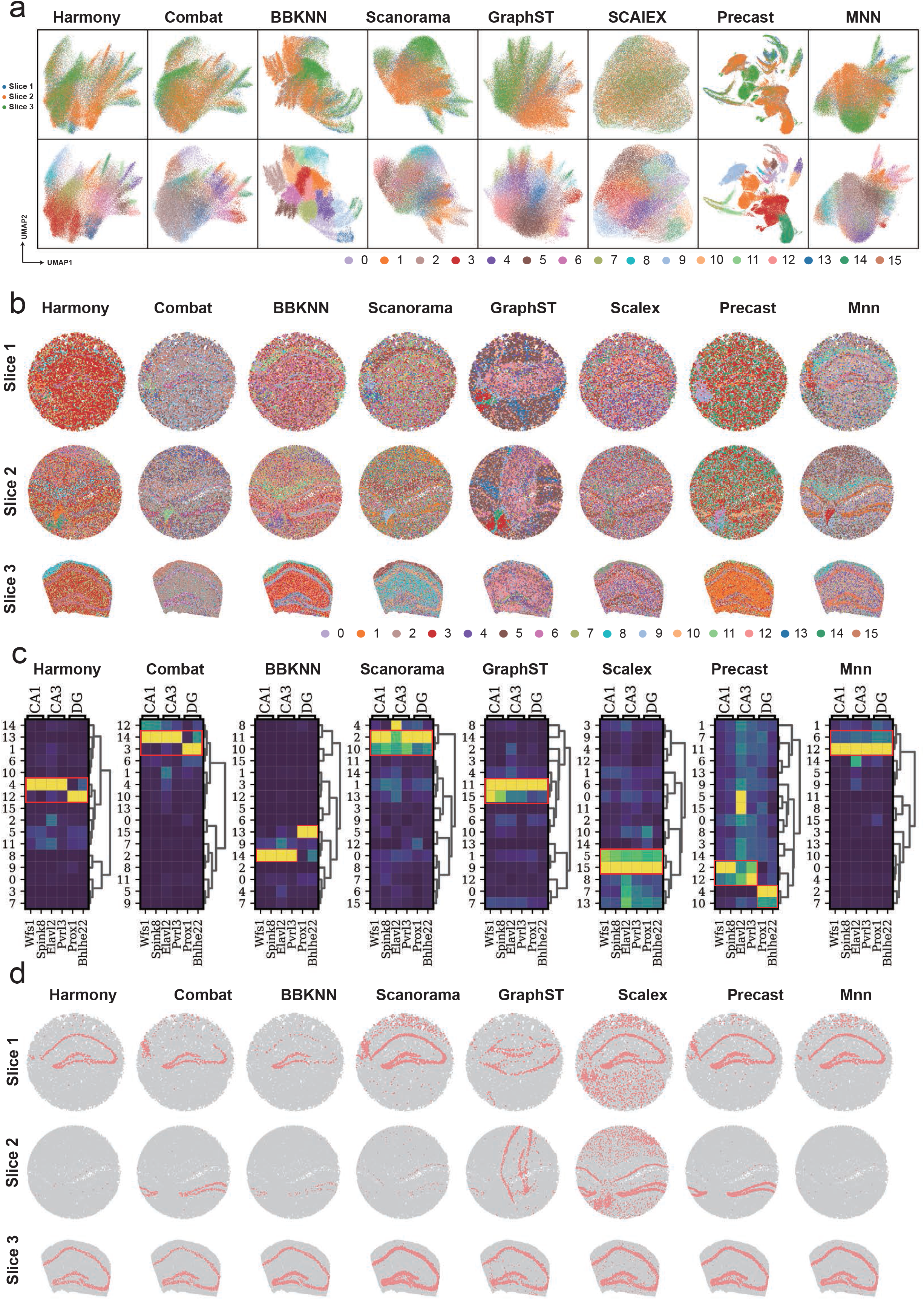
Integration results of three mouse hippocampus slices from the control methods, related to Figure 4. **a)**. UMAP plots for the joint clustering result from the control methods, coloured by slices (top panel) and cluster labels (bottom panel). **b)**. Spatial visualization of the joint clustering results from the control methods on the three slices. **c)**. Expression heatmaps of marker genes for the CA1, CA3, and DG regions in joint clusters from spatiAlign and the control methods. Clusters with high expression specificity are highlighted by red boxes. **d)**. Spatial visualization of the hippocampus-related regions on three slices identified by the control methods.

**Supplementary Fig. S6.**
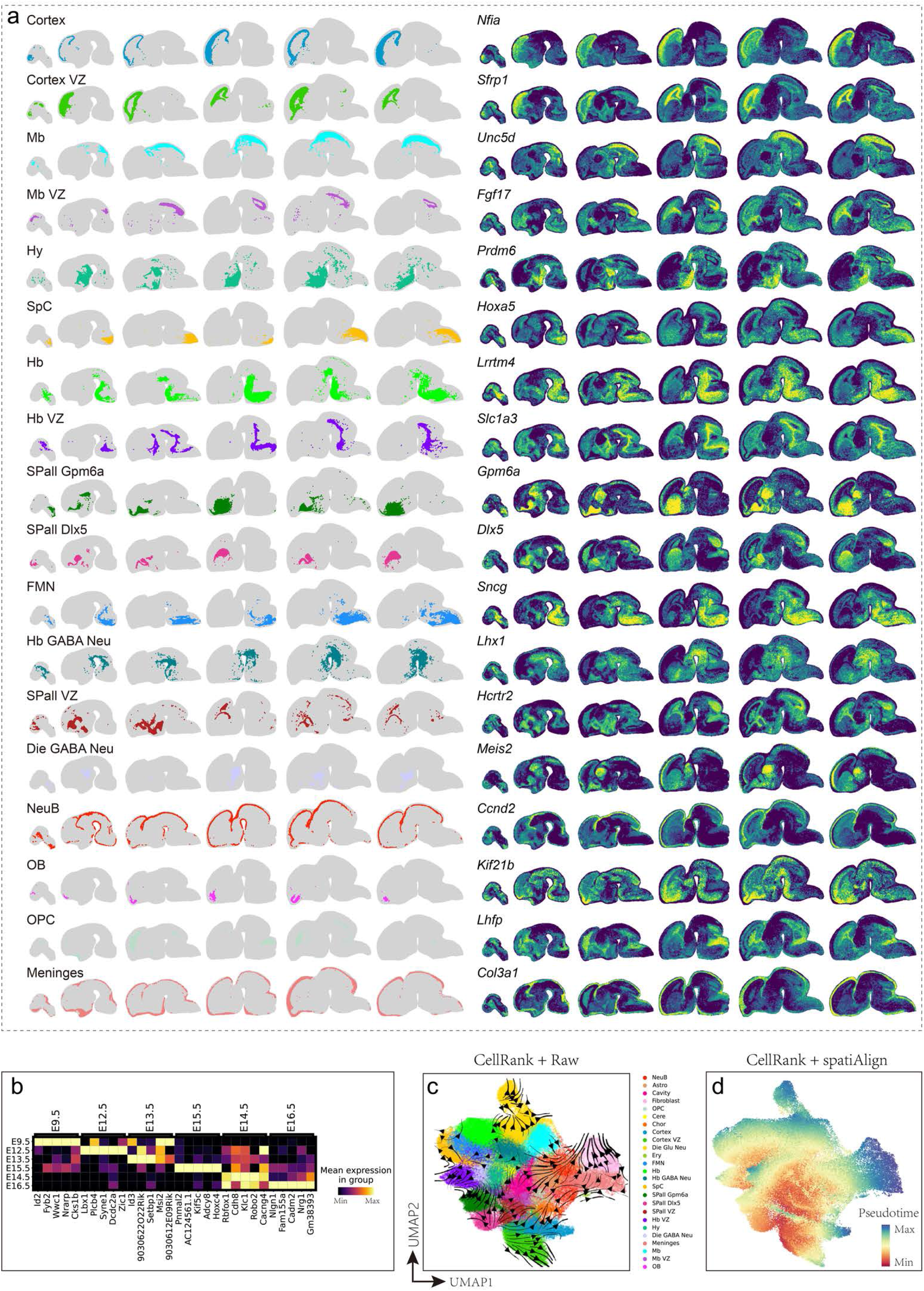
Application to time-series mouse embryonic brain, related to Figure 5. **a)**. Spatial visualization of the labelled clusters and the corresponding marker genes. **b)**. Expression heatmap of the top five differentially expressed genes from E9.5 to E16.5. **c)**. CellRank trajectory of cell types reconstructed using the raw expression counts. **d)**. Estimated pseudotime scores by spatiAlign-corrected gene expression matrices.

