## supplementary table for "spatiAlign: An Unsupervised Contrastive Learning Model for Data Integration of Spatially Resolved Transcriptomics"

**Supplyment Table 1:**

| | Sample name | Reference |
|---|---|---|
| **Slice 1** | **Puck_190921_21** | **[1]** |
| **Slice 2** | **Puck_180413_7** | **[2]** |
| **Slice 3** | **puckCropped_hippocampus** | **[3]** |


1. Rodriques SG, Stickels RR, Goeva A *et al*: **Slide-seq: A scalable technology for measuring genome-wide expression at high spatial resolution**. *Science* 2019, **363**(6434):1463-1467.
2. Wang I-H, Murray E, Andrews G *et al*: **Spatial transcriptomic reconstruction of the mouse olfactory glomerular map suggests principles of odor processing**. *Nature neuroscience* 2022, **25**(4):484-492.
3. Cable DM, Murray E, Zou LS *et al*: **Robust decomposition of cell type mixtures in spatial transcriptomics**. *Nature Biotechnology* 2022, **40**(4):517-526.
